# Assessing Aquatic Avian Biodiversity along Hydropower Dam Gradient: Understanding Impacts of Kurichhu Dam-Induced River Fragmentation in Kurrichu River Basin

**DOI:** 10.1101/2024.09.18.613804

**Authors:** Tshering Nidup, Yonten Dorji

## Abstract

Humans are the dominant ecological and evolutionary force on the planet today, transforming habitats, polluting environments, and changing climates. Obstruction to the passage of flowing rivers has a significant environmental impact on fish. It causes rippling effects on wildlife, food web, and habitat throughout the river ecosystem. However, little is known about how these fragmented habitats impact aquatic avifaunal diversity and composition along the stretch of the river. The Kurichhu hydropower dam has disconnected the continuity of upper and lower portions of the Kurichhu River for over 14 years. To investigate this, avian community data were collected at 51 point-count stations, with 17 plots each: in R (Reservoir), AR (Above Reservoir), and BD (Below Dam) along the Kurichhu hydropower dam gradient. In total, 587 individual birds of 33 species distributed under 14 families were encountered. Shannon-wiener diversity among these three sites was higher for the R pool (*H′* = 3.04) than AR (*H′* = 2.74), and the lowest was in BD (*H′* = 2.44). Menhinicks richness index was higher for AR (*D* = 2.40) than the R pool (*D =* 1.95) and lowest found in BD (*D =* 1.73). This study recorded two species of birds in vulnerable and another two species of the order *charadriformes* in the near-threatened category. Our findings found that habitats along Kurichhu, around Gyelposhing area, are an important stopover and staging area for migratory waterbird species. In light of these findings, addressing the ecological ramifications of river fragmentation on avian diversity is crucial. The marked variation in species richness and diversity across the river segments highlights the pressing need for targeted conservation strategies. To mitigate adverse impacts, we recommend implementing ecological flow regimes and creating wildlife corridors to reconnect fragmented habitats, enhancing habitat quality and supporting avian biodiversity.

## 1. Introduction

Lord and Norton (1990) defined fragmentation as the disruption of ecosystem continuity to a large extent, and the concept of fragmentation has been colored by its importance in conservation issues at the landscape scale. Thus, the direct effects of habitat transformation provide biologists with the opportunity to investigate the impacts of habitat size, quality, habitat isolation, and the effects of edges and disturbances on gene flow, populations, species, communities, and ecosystems (Fukami and Wardle, 2005; Laurance, 2008). Humans are the dominant ecological and evolutionary force on the planet today, transforming habitats, polluting environments, changing climates, introducing new species, and causing other species to decline in number or go extinct (Pelletier and Coltman, 2018). As such, fragmented habitats are frequently used to establish small population sizes and isolation that negatively influence gene flow and increase inbreeding depression (Morgan, 1999; Jump and Penuelas, 2006).

According to IUCN (2014), there are 10,425 species of birds. Of these, 1,373 species are considered threatened with extinction, 959 species are near threatened, 7,886 species are considered to be of least concern, 62 species lack the data to determine their status, and 145 species are recorded as extinct. Asian continent harbors more globally threatened water bird species than any other region of the world (Vié *et al*., 2009). Bhutan is now home to 699 species of birds (Wangdi, 2015), with 18 species of threatened status (Birdlife International, 2015). Global extinction rates are currently 100 – 1000 times greater than background levels (Dirzo and Raven, 2003; Vié *et al*., 2009).

The alarming decline of global biodiversity, particularly among vertebrate species, is a pressing concern that has seen a staggering 52% decrease in populations from 1970 to 2010, largely attributed to habitat loss, exploitation, and climate change (WWF Living Planet Report, 2014; Amhed et al. 2023). In the Himalayan region, these issues are exacerbated by anthropogenic activities driven by an increasing demand for natural resources due to population growth and economic development (Amhed et al., 2023; Waiba and Dorji, 2024).

The construction of dams, such as the Kurichu hydropower plant, poses significant ecological challenges, obstructing river flow and disrupting the delicate balance of aquatic ecosystems. Dams disrupt a river’s natural course and flow, replace turbulent river sections with still water bodies, impact flow and temperature regimes and sediment transport, alter water temperatures in the stream, redirect river channels, transform floodplains, and disrupt river continuity (Liermann *et al*., 2012; Fearnside, 2013). According to the International Rivers (2015) report, about 76 locations in Bhutan are identified for hydropower projects to be built with a potential capacity of 23,760 MW.

This study aims to fill a critical knowledge gap regarding the impacts of dam construction on avifaunal diversity in this region, specifically assessing the distribution and community composition of aquatic birds both upstream and downstream of the dam. To achieve this aim, our research will focus on two primary objectives:

1. First, we will compare species richness, diversity, and abundance of aquatic avifauna in areas above and below the Kurichu dam. This comparative analysis will provide insights into how dam-induced alterations in habitat structure affect bird populations.
2. Second, we will establish baseline data on the winter community composition of aquatic birds in this region. Understanding these dynamics is essential for evaluating the ecological consequences of dam construction on avian diversity and developing informed conservation strategies.

The ecological ramifications of obstructing river systems are profound, extending beyond immediate impacts on fish populations to affect entire food webs and wildlife habitats. Dams can lead to altered sediment transport, changes in water temperature and chemistry, and disruptions in migratory patterns for various species. These changes can have cascading effects throughout the ecosystem, ultimately impacting not only avifauna but also the broader biodiversity that relies on healthy riverine environments.

This research represents a pioneering effort to assess the ecological impacts of Hydropower dam on aquatic bird communities. By comparing avifaunal diversity above and below the Kurichu dam and establishing baseline data for future studies, we hope to contribute valuable knowledge to inform conservation efforts and sustainable resource management practices.

## 2. Materials and Methods

### 2.1 Study area

The study was conducted along the Kurichhu River, assessing both the above and below hydropower dam during winter (Fig. 1). The Kurichhu Hydropower Plant is located at Gyalpozhing, Mongar, on the Kurichhu River in Eastern Bhutan, with a dam height of 55 m (from its deepest foundation), crest length of 285 m, and a surface powerhouse located at the toe of the dam. The Project has an installed capacity of 60 MW consisting of four units of 15 MW each and a mean annual energy generation capacity of 400 million units (MU) (Druk Green Power Corporation [DGPC], 2015). Geographically, the hydropower plant was located at a longitude of 91° 21’E and latitude of 27° 13’N with an altitude range of 531 masl (DGPC, 2015).

**Figure 1.**
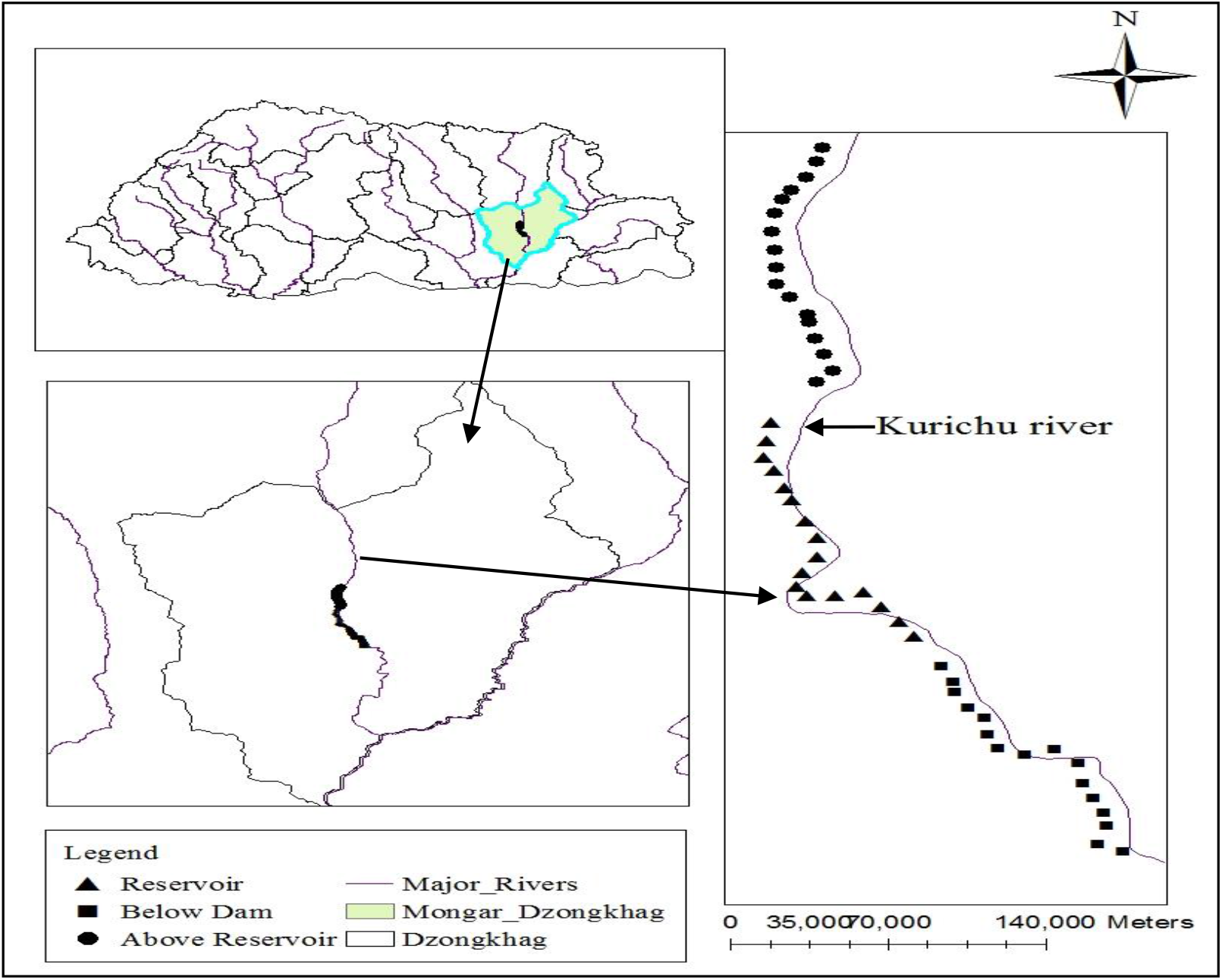
Location of the study area, showing the detailed research plots along the Kurichhu hydropower dam

### 2.2 Stratification of the study area and sampling design

The study area was stratified into three clusters: a) distance measured from the dam to the length of reservoir pool formed by the dam towards the North; b) this same length of distance was replicated downstream of the dam Southwards, and c) equal length of distance was again replicated above the reservoir pool towards further North. Fixed-radius plots were laid out along these three replicates with 300 m in between these plots to avoid repeated counting of birds; this followed the published methodology used in Bibby *et al*. (2000). Bird survey was done using direct count methods. A total of 51 sampling points covering a distance of 15300 m were surveyed and enumerated following the point count sampling method developed by Bibby *et al*. (2000)

### 2.3 Sampling method

The assessment of birds along the Kurichhu hydropower dam was done in the winter season. The sample plots were visited twice a week for enumeration. Data collection was carried out for 5 hours a day, from 0630 hrs. to 1030 hrs. in the morning and from 1630 hrs. to 1730 hrs. in the afternoon, when the activities of birds were prominent, which commenced about 30 min after dawn and continued to mid-morning. Observers are rotated among stations to distribute potential differences in observer ability. We also rotated the order of stations visited according to the time of day. During each visit, all birds seen or heard within 50 (0-15; 15-30; 30-50) meters radius of 15 min duration at each point were recorded in the datasheet. To minimize disturbance during the count, a waiting period of 3 to 5 min before counting was applied, as followed by Bibby *et al*. (2000). Counting was accomplished for a fixed period of 3-10 min, depending upon how conspicuous the birds were as done in the study by Zakaria and Rapar, (2013).

Identification of birds was confirmed using the book on Birds of Bhutan by Inskipp *et al*. (2011) and Birds of Indian Subcontinent by Grimmet, *et al*. (1999). Birds sighted during the survey were categorized based on their migratory nature, including Passage Migrant (PM), Resident (R), Altitudinal Migrant (AM), Winter Visitor (WV), and Summer Visitor (SV), following the well-developed methods by Inskipp *et al*., (2011). The nomenclature, taxonomic listing, and conservation status of the avifauna were determined based on the IUCN Red List of Threatened Species (2014).

### 2.4 Data analysis

The data collected during the study period were compiled and analyzed using Microsoft Excel and R software. Since the data point per variable was less than 30, non-parametric tests were performed. The following formulae were used to calculate species diversity, richness, relative abundance, and composition. The variables used for the test include the Shannon diversity index and the Menhinick richness index.

A. Shannon Wiener diversity index (*H′*) *H′* = -∑ P_i_ × ln(P_i_), where *H′* = index of species diversity, P_i_ = Proportion of total sample belonging to the *i*th species, ln = natural logarithm.
B. Menhinick Species richness (*D*) index = the ratio of the number of taxa to the square root of sample size.
C. Relative abundance *(RA)* expressed in (%) = the ratio of a number of particular species to the total detected species.
D. Jaccard similarity index, 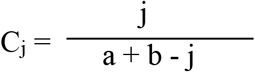 where, j = the number of species found at both sites A and B, a = the number of species in site A, and b = the number of species found in site B.

These indices are designed to equal 1 where the species from the two sites are the same and 0 if the sites have no species in common.

(E) Renkonen index (P) = ∑ minimum 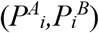Where, *P*^*A*^_*i*_, is the percentage of species *i* in sample A *P*^*B*^_*i*_, is the percentage of species *i* in sample B S is the number of species found in either sample. With no overlap between samples, the index equals 0; with complete similarity, the Renkonen index equals 100%.

## 3. Results and discussion

### 3.1 Bird species composition and relative abundance

A total of 587 birds distributed under 14 families were encountered along the Kurichh hydropower dam over the study period (Table 2). The birds that were detected in the three sites sampled were from the families *Accipitridae, Alcedinidae, Anatidae, Ardeidae, Charadriidae, Cinclidae, Motacillidae, Muscicapidae, Pandionidae, Phalacrocoracidae, Podicipedidae, Rallidae, Scolopacidae*, and *Turdidae*. Out of 14 families recorded, *Anatidae* had the highest number of bird species (eight species), followed by *Muscicapidae* (five species), *Charadriidae, Alcedinidae, Ardeidae* (three species) each. Family *Rallidae* and *Motacillidae* had (two species) each, while *Scolopacidae, Phalacrocoracidae, Accipitridae, Turdidae, Cinclidae, Pandionidae* and *Podicipedidae* had single species each. Vyas *et al*. (2012) noted that family *Anatidae* was most dominant family during winter month from Barna reservoir, represented by 16 species. The maximum composition of species was observed in Reservoir pool (RP) (*n* = 26), followed by Above Reservoir (AR) (*n* = 25). The least was found Downstream of Dam (DD) (*n* = 17).

The highest number of species in *Anatidae* family may be due to the presence of passage migratory species inhabiting different habitats along Kurichhu River. And also pool created by dam and other similar habitats for dabbling ducks [e.g., Common Pochard (*Aythya ferina* Linnaeus, 1758), Red-crested Pochard (*Netta rufina* Pallas, 1773), Mallard (*Anas platyrhynchos* Linnaeus, 1758), Common Merganser (*Mergus merganser* Linnaeus, 1758), and Great crested grebe (*Podiceps cristatus* Linnaeus, 1758)]. Most of the birds observed during this study were resident and migratory species. Lameed (2011) reported that the species that are winter migrants use wetlands for resting till their home range conditions become favorable. The most frequently detected species were Great Cormorant (*n*_*J*_ = 45; *n*_*F*_ = 40) with relative abundance (RA) of 16.25% in January and 12.90% in February months, Plumbeous Water Redstart (*n*_*J*_ *=* 38; *n*_*F*_ = 39) with RA 13.72% in January and 12.58% in February. White-capped Water Redstart (*n*_*J*_ *=* 37; *n*_*F*_ = 33) with 13.36% in January and 10.65% in February. Where *n*_*J*_ and *n*_*F*_ are the number of birds detected in January and February months, respectively. Among these three sample sites, a higher number of birds were observed in the Above Reservoir (109 individuals, 35.39%) than in the Reservoir (102 individuals, 33.11%) and lower in Downstream Dam (97 individuals, 31.5%). The difference of two percent margin may be attributed to having greater resources such as food and nesting sites and the ability to support more birds (Soka *et al*., 2013) in such habitats.

### 3.2 Spatial distribution and composition of birds

The overall diversity of birds and species richness of the three samples are shown in Table 1 and Figure 3 above, where the highest Shannon diversity index was observed in Reservoir (*H* ^′^ = 3.04). The lowest value below the dam (*H* ^′^= 2.44) and the Menhinicks richness index ranges from *D* = 2.39 to *D* = 1.73 (Fig. 2). Shannon diversity (*H* ′) among these three sample sites were statistically significant, with *H* (2) *=* 7.627, *p* = 0.02; however, Menhinicks richness among three sample sites did not show significant values (*H* (2) = 4.867, *p* > 0.05). The habitats of these three samples vary differently from each other. Habitats above the dam comprise a reservoir pool created by Kurichhu dam. This change in habitat to pool helps in the assemblage of species from *Anatidae* family en route on their migration and other edge-inhabiting species. The high diversity in reservoirs is supported in a study by Fruget (1992), ascertaining that waterbird assemblages respond strongly to dam-induced habitat changes - populations often increase on reservoirs in response to development of new habitats and food resources and are indicators of ecosystem change. Jaccard similarity index (C_j_) and Renkonen similarity index (p) were 0.645 and 51.4%, respectively, between the AR (Above Reservoir) and R (Reservoir).

**Table 1.**
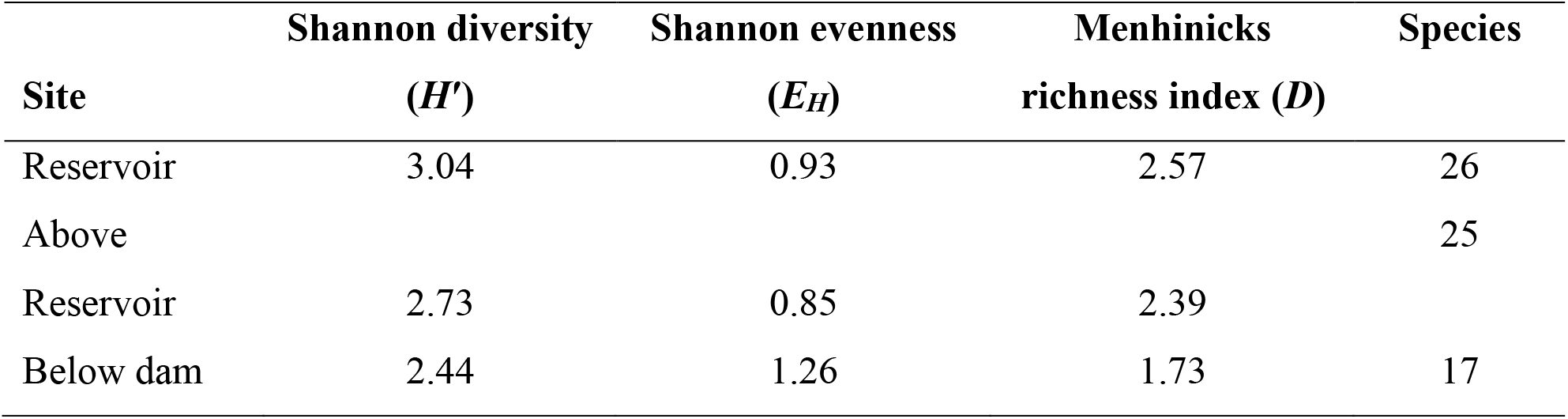
Diversity, evenness, and species richness of the study area.

| Site | Shannon diversity<br>($H'$) | Shannon evenness<br>($E_H$) | Menhinicks<br>richness index ($D$) | Species |
| --- | --- | --- | --- | --- |
| Reservoir | 3.04 | 0.93 | 2.57 | 26 |
| Above |  |  |  | 25 |
| Reservoir | 2.73 | 0.85 | 2.39 |  |
| Below dam | 2.44 | 1.26 | 1.73 | 17 |

**Table 2.** Checklist of birds recorded during the study.

| Sl. No. | Order/family | Common name | Scientific name | MS | IUCN 2015 |
| --- | --- | --- | --- | --- | --- |
| 1 | *Anseriformes/ Anatidae* | Common Merganser | *Mergus merganser* | WV | LC |
| 2 |  | Common Teal | *Anas crecca* | WV/PM | LC |
| 3 |  | Common Pochard | *Aythya ferina* | PM | VN |
| 4 |  | Eurasian Wigeon | *Mareca penelope* | WV/PM | LC |
| 9 | *Coraciiformes/ Alcedinidae* | Common Kingfisher | *Alcedo atthis* | R | LC |
| 10 |  | Crested Kingfisher | *Megaceryle lugubris* | R | LC |
| 11 |  | White-throated Kingfisher | *Halcyon smyrnensis* | R | LC |
| 12 | *Pelecaniformes/ Ardeidae* | Black-crowned Night Heron | *Nycticorax nycticorax* | R/SV | LC |
| 13 |  | Cattle Egret | *Bubulcus ibis* | R | LC |
| 14 |  | Little Heron | *Butorides striatus* | R | LC |
| 15 | *Charadriiformes/ Charadriidae* | Red-wattled Lapwing | *Vanellus indicus* | R | LC |
| 16 |  | River Lapwing | *Vanellus duvaucelii* | R | NT |
| 17 |  | Northern Lapwing | *Vanellus vanellus* | PM | NT |
| 18 | *Scolopacidae* | Common Sandpiper | *Actitis hypoleucos* | WV/PM | LC |
|  | *Suliformes/* |  |  |  |  |
| 19 | *Phalacrocoracidae* | Great Cormorant | *Phalacrocorax carbo* | WV/PM | LC |
|  | *Accipitriformes/* |  |  |  |  |
| 20 | *Accipitridae* | Pallas's Fish Eagle | *Haliaeetus leucoryphus* | R/PM | VN |
| 21 | *Pandionidae* | Osprey | *Pandion haliaetus* | WV/PM | LC |
|  | *Passeriformes/* |  |  |  |  |
| 22 | *Muscicapidae* | Hodgson's Redstart | *Phoenicurus hodgsoni* | WV | LC |
| 23 |  | Little Forktail | *Enicurus scouleri* | AM | LC |
| 24 |  | Plumbeous Water Redstart | *Rhyacornis fuliginosus* | AM | LC |
| 25 |  | Slaty-backed Forktail | *Enicurus schistaceus* | AM | LC |
| 26 |  | White-capped Water Redstart | *Chaimarrornis leucocephalus* | AM | LC |
| 27 | *Motacillidae* | Grey Wagtail | *Motacilla cinerea* | AM | LC |
| 28 |  | White Wagtail | *Motacilla alba* | AM/W<br>V | LC |
| 29 | *Turdidae* | Blue Whistling Thrush | *Myophonus caeruleus* | AM | LC |
| 30 | *Cinclidae* | Brown Dipper | *Cinclus pallasii* | AM | LC |
|  | *Podicipediformes/* |  |  |  |  |
| 31 | *Podicipedidae* | Great Crested Grebe | *Podiceps cristatus* | PM | LC |
|  | *Gruiformes* |  |  |  |  |
| 32 | *Rallidae* | Common Moorhen | *Gallinula chloropus* |  | LC |
| 33 |  | Ruddy-breasted crake | *Zapornia fusca* | R/AM | LC |

**Figure 2.**
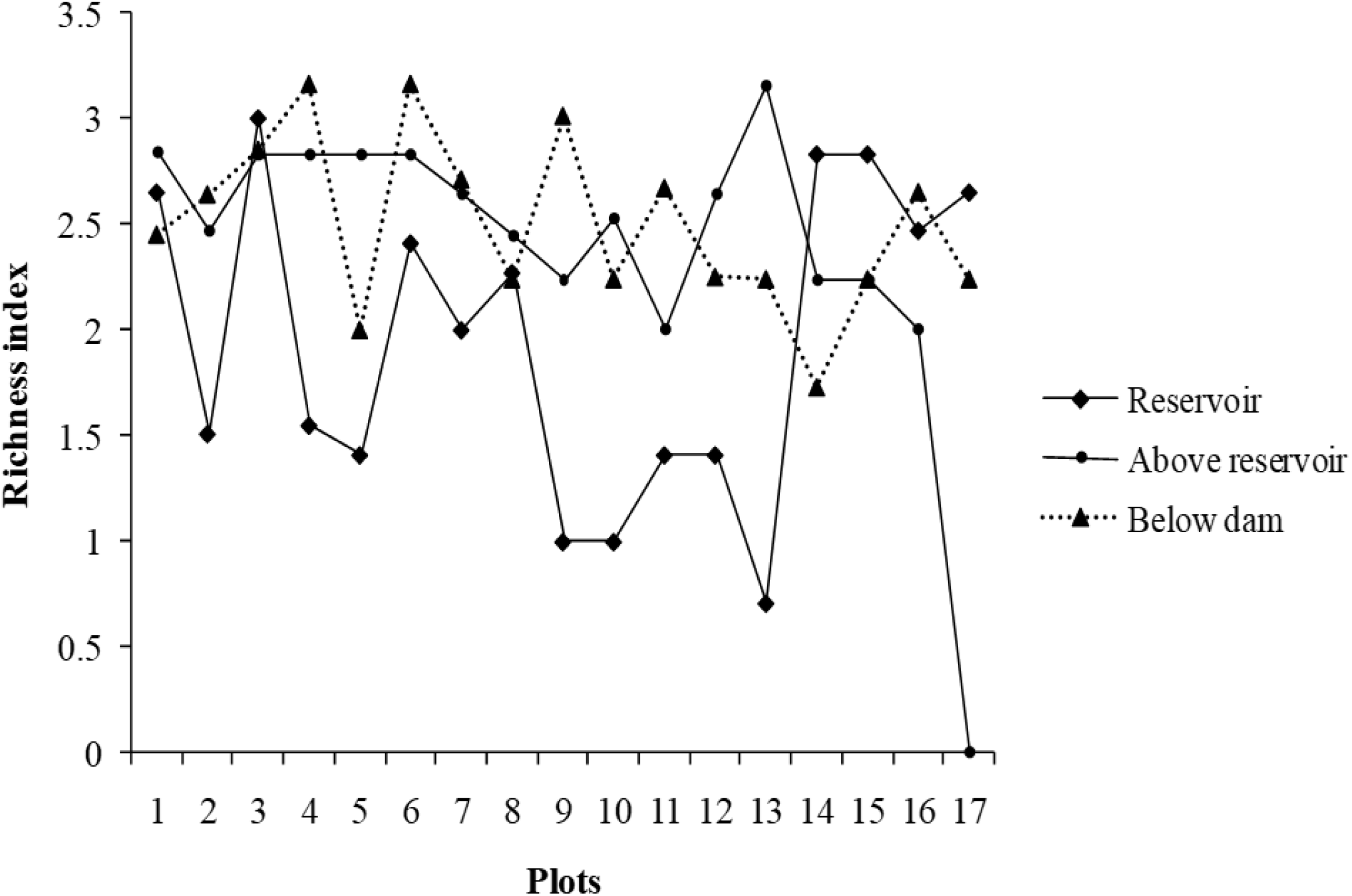
Menhinick richness index of the study area (R, AR, BD)

**Figure 3.**
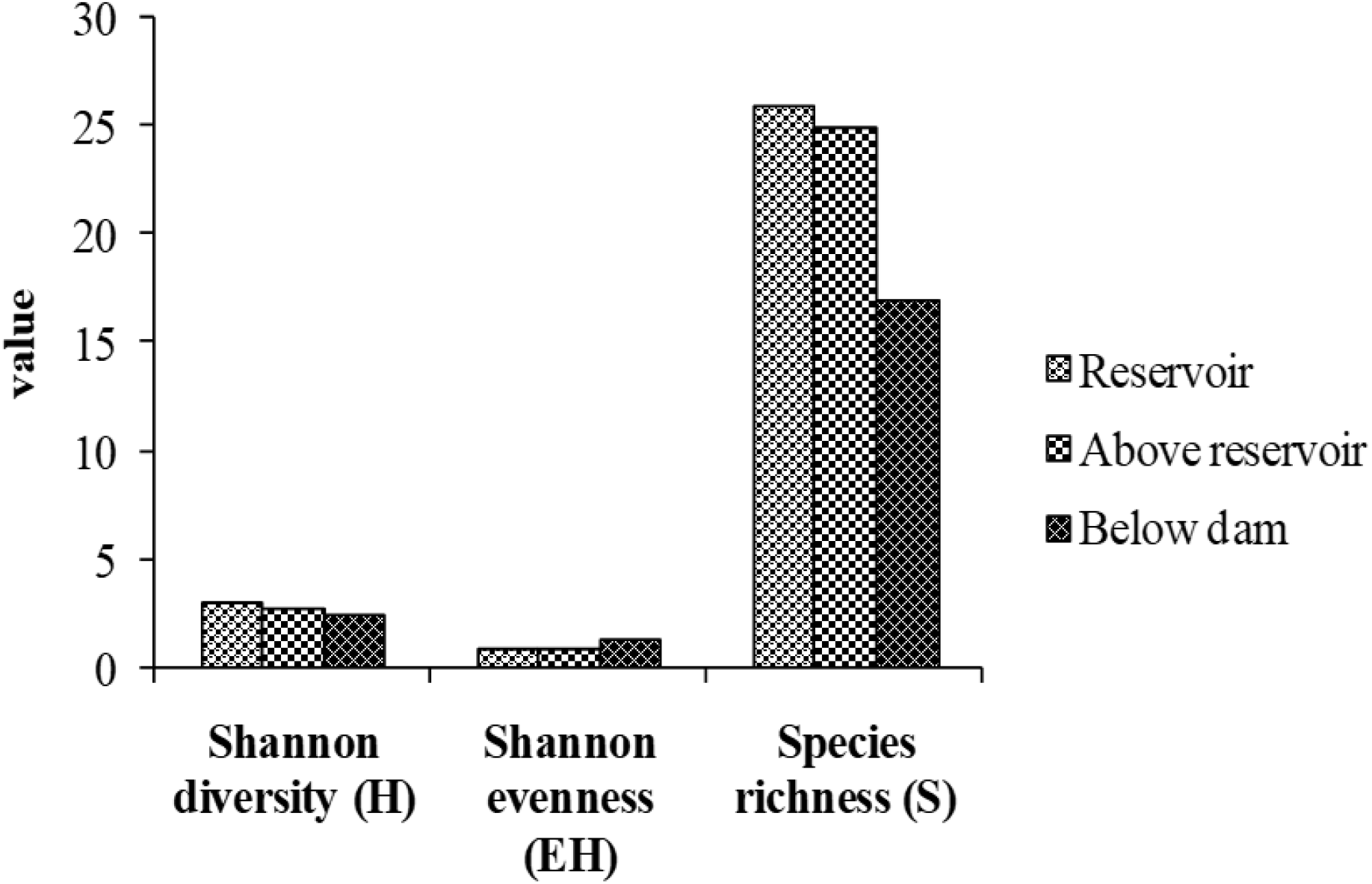
Diversity, Evenness, and Species richness in the study area ( R, AR, BD)

Similarly, the Jaccard index and Rekonen index between the R (Reservoir) and BD (Below Dam) were 0.43 and 46.8%, respectively. This implies that almost half of the species are similar between R to AR and R to BD. However, the Jaccard similarity index between the above Reservoir and below dam is 0.62, and the Renkonen index is 78%. Habitats of both AR and BD comprise of fast flowing and running rivers, where species such as Plumbeous Water Redstart, White-capped Water Redstart, Little Forktail, White Wagtail and Blue Whistling Thrush of *Muscicapidae* family were widespread, similar to a study by Dorji (2015). Great Cormorants, River Lapwing and Common Sandpiper were also observed in the entire site, along with a few species of *Anatidae* family, such as Common Merganser.

The study also records globally threatened species such as the Common Pochard (*Aythya ferina* Linnaeus, 1758) and Pallas’s Fish Eagle (*Haliaeetus leucoryphus* Pallas, 1771) in the vulnerable (VU) category. River Lapwing (*Vanellus duvaucelii* Lesson, 1826) and Northern Lapwing (*Vanellus vanellus* Linnaeus, 1758) are categorized as near threatened (NT).

## 4. Conclusion and Recommendation

The mosaic of habitats along Kurichhu Dam makes it a unique avifaunal refuge for many migratory and residential waterbirds. Hence, the place around Gyelposhing is a vital staging and resting place for birds en route to migration. The area also provides a breeding habitat for Pallas’s Fish Eagle and River Lapwing, categorized as threatened species by IUCN.

This study provides clear evidence of the profound ecological effects that river fragmentation, caused by the Kurichhu hydropower dam, has on avian diversity and composition. The significant variations in species richness and diversity across the reservoir, above, and below dam sites indicate that habitat fragmentation disrupts natural ecological processes, adversely affecting avifaunal communities. The higher diversity in the reservoir area compared to the upstream and downstream regions suggests that habitat alteration, particularly the creation of standing water in the reservoir, creates conditions more favorable for certain bird species, likely those adapted to lentic systems. Meanwhile, the lower diversity and richness below the dam (BD) reflects the detrimental impact of disrupted river flow on downstream habitat quality, which in turn affects species dependent on lotic environments.

The presence of vulnerable and near-threatened bird species, mainly migratory waterbirds and species of the order Charadriiformes, underscores the global ecological significance of the Kurichhu riverine habitats. The altered flow regimes have likely disrupted the ecological integrity of this region, with implications not only for local biodiversity but also for broader ecological networks, including the migratory routes that rely on these habitats.

To address the observed ecological impacts of river fragmentation, we strongly recommend the implementation of ecological flow regimes to restore natural flow variability and improve downstream habitat conditions. These regimes can help maintain the ecological connectivity essential for avian and aquatic biodiversity by mimicking the natural hydrological patterns. Furthermore, establishing wildlife corridors along the river would reconnect fragmented habitats, facilitating the movement and dispersal of species, particularly migratory birds, and mitigating the adverse effects of habitat isolation.

In addition, long-term monitoring of avian diversity and habitat conditions is crucial to assess the effectiveness of these conservation strategies. We recommend expanding this research to include other taxa, such as fish and invertebrates, to understand better how river fragmentation impacts biodiversity at multiple trophic levels. This broader ecological perspective will inform the development of more integrated and sustainable conservation policies that balance hydropower development with biodiversity conservation. Ultimately, this data-driven approach will enhance our understanding of avian dynamics and enable us to implement targeted and effective conservation strategies, supporting the preservation of our avian biodiversity and the health of our ecosystems.

## Acknowledgment

We thank the Druk Green Power Corporation Limited (DGPC) for allowing us to conduct this study. We acknowledge the College of Natural Resources, Royal University of Bhutan, for the technical and necessary logistic support and for providing a conducive environment during this study.

